# Schizophrenia Risk Alleles Often Affect the Expression of Many Genes and Each Gene May Have a Different Effect on The Risk; A Mediation Analysis

**DOI:** 10.1101/2020.01.27.904680

**Authors:** Xi Peng, Joel S. Bader, Dimitrios Avramopoulos

## Abstract

Variants identified by genome-wide association studies (GWAS) are often expression quantitative trait loci (eQTLs), suggesting they are proxies or are themselves regulatory. Across many datasets analyses show that variants often affect multiple genes. Lacking data on many tissue types, developmental time points and homogeneous cell types, the extent of this one-to-many relationship is underestimated. This raises questions on whether a disease eQTL target gene explains the genetic association or is a by-stander and puts into question the direction of expression effect of on the risk, since the many variant - regulated genes may have opposing effects, imperfectly balancing each other. We used two brain gene expression datasets (CommonMind and BrainSeq) for mediation analysis of schizophrenia-associated variants. We confirm that eQTL target genes often mediate risk but the direction in which expression affects risk is often different from that in which the risk allele changes expression. Of 38 mediator genes significant in both datasets 33 showed consistent mediation direction (Chi^2^ test P=6*10^−6^). One might expect that the expression would correlate with the risk allele in the same direction it correlates with disease. For 15 of these 33 (45%), however, the expression change associated with the risk allele was protective, suggesting the likely presence of other target genes with overriding effects. Our results identify specific risk mediating genes and suggest caution in interpreting the biological consequences of targeted modifications of gene expression, as not all eQTL targets may be relevant to disease while those that are, might have different than expected directions.

## INTRODUCTION

Schizophrenia (SZ) is a common and disabling mental disorder with a point prevalence of 0.5%, an onset in late adolescence or early adulthood and a lifelong course with significant disability [Messias and others 2007]. It has high heritability, among the highest in psychiatric disorders, consistently estimated around 80% [Cardno and others 1999; Kety 1987]. Although recognized diagnostically as a single disorder, SZ has a highly heterogeneous phenotype with symptoms that range from prominent delusions, hallucinations, agitation, and erratic behavior to lack of interest and motivation, apathy and disorganization. The response of patients to different treatments is also highly variable. While some of this heterogeneity might be the result of environmental effects, a large proportion of the variability likely reflects the underlying genetic heterogeneity and the compromise of different biological processes that alone or in combinations lead to the phenotype we call “SZ”. Once we understand the links between genetic variation and biological processes, genetic testing might predict each patient’s response to treatment or liability to environmental exposures. This would be a big step forward in personalized prevention and treatment, a benefit for the patient and the society.

Over the last few years, thanks to large collaborative genome-wide association studies (GWAS) such as by the Psychiatric Genomics Consortium (PGC: https://pgc.unc.edu/), SZ has become the psychiatric disorder with the largest number of genetic variants robustly shown to contribute to risk. The number of SZ loci has steadily increased from 5 to 22 and currently over 100 [Consortium 2011; Consortium 2014; Pardinas and others 2018; Ripke and others 2013]. As observed with GWAS-identified loci across other complex disorders the SZ-associated variants are most often located in non-coding sequences and 40% of the time their haplotype blocks do not include coding exons [Hindorff and others 2009; Manolio and others 2009; Visel and others 2009]. These variants are presumed to be regulatory, which is supported by their concentration in open chromatin DNA as marked by deoxyribonuclease I (DNase I) hypersensitive sites (DHSs) [Maurano and others 2012] and by studies of Quantitative Trait Loci (eQTLs) [Cookson and others 2009; Gilad and others 2008; Hindorff and others 2009; Majewski and Pastinen 2011; Schaub and others 2012]. Regulatory sequences, however, can be far from their target gene, so it is difficult to assign a specific gene or genes to each variant solely by location. In fact, it has been shown using chromatin interaction data that in most cases the nearest gene to the variant is not the one affected by it [Maurano and others 2012]. Further, regulatory sequences often regulate more than one gene as shown by interactions with multiple promoters [Akerborg and others 2019] and consistently observed in eQTL databases (GTEx: gtexportal.org, Commonmind: www.nimhgenetics.org/resources/commonmind and BrainSeq: eqtl.brainseq.org). Further, as eQTL discovery depends on the studied cell type, tissue and developmental time point and current studies are far from covering all these possibilities, it is likely that there remain undiscovered variant-gene correlations and that many more eQTLs might regulate multiple genes rather than one gene. In fact, because studies of eQTLs specific to development are rare [O’Brien and others 2018] and usually done in bulk tissue, many eQTLs likely remain unknown. These missing eQTLs are also the most likely to be of importance for a developmental brain disorder like SZ. With multiple genes regulated by the same variant it remains unknown whether it is the gene(s) we know off that mediate the increase in disease risk and whether it is only one or more target genes of the same variant. Figure 1 illustrates how many genes regulated by one risk allele may all exert effects on the risk, and these effects may be different with the final effect of the allele being the sum of them all. The consequence of this is that if a disease risk allele increases the expression of the gene it does not necessarily follow that increased expression translates to higher risk. This is of great importance as disease-modeling studies often perturb gene expression to study the outcome and understand the biology of disease [Das and others 2020; Hill and others 2012; Schrode and others 2019; Yang and others 2018]. It is therefore necessary to seek formal evidence that a specific gene’s expression mediates disease risk and in which direction, if we want to have an accurate list of the genes and a correct understanding of their role in disease.

**Figure 1:**
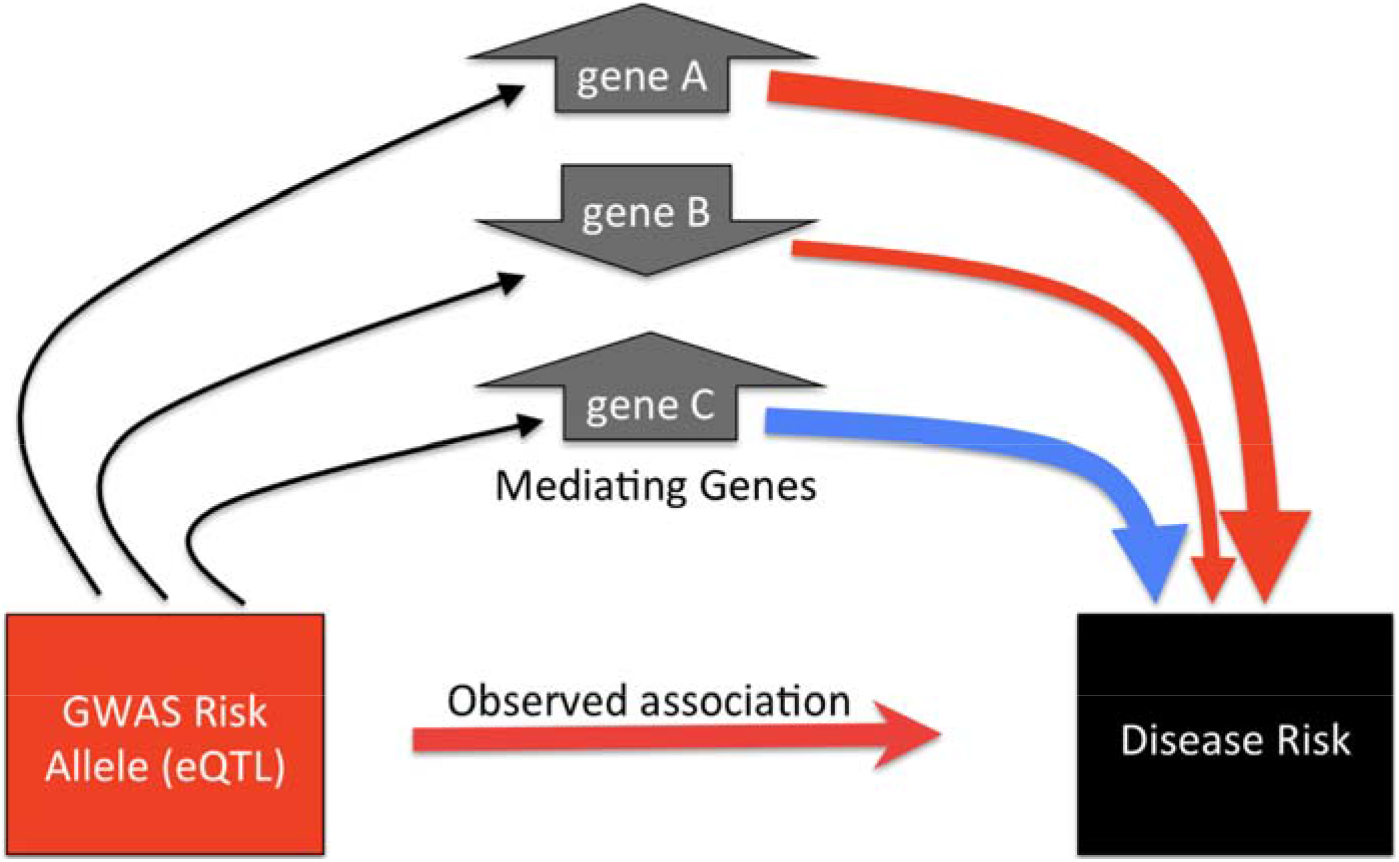
Mediation through multiple genes. The risk allele changes the expression of multiple genes up- or downward (indicated by the arrow box around each gene). This modification may increase risk (curved red arrows, width reflects effect size) or decrease it (blue arrow). The association of the risk allele with higher risk (straight red arrow) reflects the sum of all these effects on risk. The mediation of some genes expression change may not follow the direction of the the risk allele’s effect, as in gene C here.

We should note that observing opposing effects from genes regulated by the same variant should probably be expected. Alleles changing the expression of a gene in a way that increases risk are subject to selective pressure, especially in schizophrenia where there is a large reduction in fecundity for patients [Power and others 2013]. If these alleles have large effects, they are unlikely to remain in the population, as demonstrated by the strong negative correlation between effect size and allele frequency [Avramopoulos 2018]. Schizophrenia alleles that influence the expression of multiple genes would be more likely to persist if these genes balance each other’s effect on risk rather than if they all contribute in the same direction

Mediation analysis is a statistical method that aims to explain the mechanism behind an association between an independent and a dependent variable (in this case genotype and disease) by the inclusion of a third mediator variable (in this case gene expression) that is hypothesized to underlie the effect. Mediation analysis tests whether a variable is in a causal sequence between the independent and dependent variables, in other words it is a mediator [MacKinnon and others 2007] (Figure 2), One of the most cited approaches and software tools have been developed by Andrew Hanes [Hayes 2018]. Importantly, once there is significant prior evidence for the association of a variable (genotype) with the outcome (disease) the “effect to be mediated”, (genotype on risk) on risk, does not need to be significant in a specific dataset in order to test for mediation [Mackinnon and Fairchild 2009; Zhao and others 2010]. This is important for variants selected from GWAS that are unlikely to consistently show statistical significance in the much smaller postmortem tissue datasets necessary to test mediation. While mediation analysis has been widely used in psychology, it has rarely been applied to test whether gene expression mediates disease risk. Formally showing mediation for disease associated variants that are eQTLs is important as it can (i) validate the assumption that a gene regulated by a variant is important for the disease; (ii) point to specific genes where this is true, as opposed to other genes regulated by the same variant; (iii) indicate the direction of the effect of the gene.

**Figure 2:**
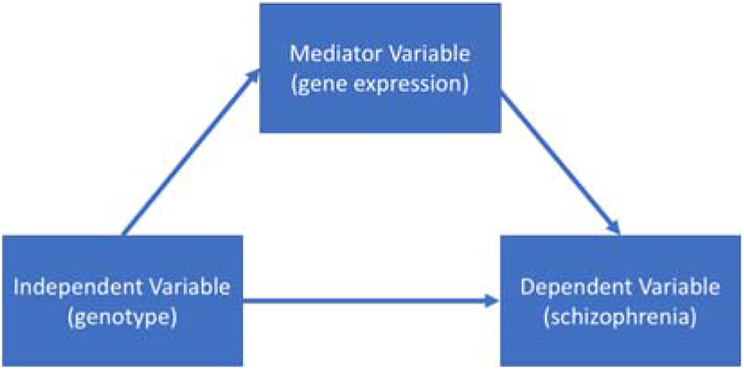
Mediation Analysis tests whether a variable is in a causal sequence between the independent and dependent variables, in this case whether gene expression is in a causal sequence between genotype and disease

Here we hypothesize that for many of the known eQTL variants that are also associated with SZ: A) The eQTL target gene frequently mediates the effect and B) the direction of effect of the risk allele on gene expression may not be the same with the direction of effect of the gene expression on the risk, presumably due to additional target genes that may not always be known. To test these hypotheses, we perform mediation analysis on two large public datasets that include both genotype and RNA-Seq data, the CommonMind Consortium (CMC) and the BrainSeq Consortium (BSC). We find evidence supporting the hypothesis of multiple target genes and report on mediating genes and their direction of effect at multiple levels of statistical confidence.

## MATERIALS AND METHODS

### Datasets

#### Psychiatric Genomics Consortium Data

Summary data from Psychiatric Genomics Consortium (PGC) [Consortium 2014] were downloaded from https://www.med.unc.edu/pgc, and a p-value threshold of 10^−6^ was used to select variants for our analysis, a relaxed threshold as this analysis is not meant to nominate SZ risk variants but rather to test our mediation and direction hypotheses. Due to the high linkage disequilibrium (LD) of the major histocompatibility complex region and instead of making decisions on the boundaries and we excluded all SNPs on chromosome 6 since this would be unlikely to influence our overall results, leaving 13,197 SNPs meeting the significance criteria. While each of these SNPs was tested for mediation, many are in high LD with each other, and the effective number of independent tests is less than then number of SNPs.

To estimate the multiple testing correction and create interpretable structure, we created groups of SNPs based on LD. We first calculated the r^2^ between pairs of SNPs using combined genotype data from CMC and BSC. We then used r^2^ ≥ 0.2 to introduce links between pairs of SNPs and defined groups as the connected components. Within groups, each SNP has at least one other SNP r^2^ ≥ 0.2; between groups, all r^2^ values are less than 0.2. This process created 219 groups (supplementary table 1). We then defined groups as the same locus if their closest SNPs were closer than 1 Mb apart.

Although SNPs within groups are correlated, they still represent multiple tests. To account for multiple testing, we performed permutations to calculate the study-wide expected number of positives under the null hypothesis and the standard deviation (SD) of this number.

#### CMC Data

We downloaded the quality controlled and normalized expression data and imputed genotype data from the CMC (https://www.synapse.org/#!Synapse:syn2759792/wiki/69613). Details on the generation of this data can be found in the group’s published work [Fromer and others 2016]. In brief this dataset contains the results of RNA sequencing data from postmortem human dorsolateral prefrontal cortex. The same individuals were genotyped using the Illumina Infinium HumanOmniExpressExome chip and imputed to the 1,000 Genomes Phase 1 reference panel. We used the RNA data already adjusted by the CMC investigators for known covariates (ancestry, sex, age at death, PMI, RIN & RIN squared, Institution, and one clustered experimental variable - see original publication [Fromer and others 2016]) and hidden covariates, generated by surrogate variable analysis, using linear regression.

We kept for analysis 258 individuals with diagnosis of “Schizophrenia” and 279 “Controls” with genotype and expression data. The cases included 214 Caucasians, 38 African Americans, 5 Hispanics, 1 Asian, and the controls 212 Caucasians, 45 African Americans, 18 Hispanics, 3 Asian, 1 Multiracial, respectively. For consistency with the BSC data, we used PLINK 1.9 to remove SNPs with genotyping rate < 0.90, minor allele frequency < 0.05, or Hardy-Weinberg P value < 10^−6^. We also removed all SNPs with multi-character allele codes, with single-character allele codes outside of A, C, T, G, or with codes missing. We then extracted the genotypes of SNPs that were also present in the PGC data with consistent alleles. Finally, we excluded all strand-ambiguous SNPs (genotypes G/C or A/T). In total 9536 SNPs and 16311 genes were included in the eQTL analysis.

#### BrainSeq Consortium Data

The pre-imputed and quality controlled genotype data and the non-quality controlled human dorsolateral prefrontal cortex expression data of the BSC [Jaffe and others 2018] were provided to us by Dr. Andrew Jaffee. Genotyping of postmortem tissue in this cohort was performed using the Illumina HumanHap650Y_V3, Human 1M-Duo_V3, and Omni5 chips, followed by imputation on the 1,000 Genomes Phase 3 reference set. As with CMC, we removed SNPs with genotyping rate < 0.90, minor allele frequency < 0.05, Hardy-Weinberg P value < 1*10^−6^. The Poly(A)+ RNA sequencing was performed by the BSC investigators using Illumina HiSeq 2000 with two hundred bp paired-end sequencing. Reads were mapped to the human genome hg19 using TopHat 2.0.4. Similar to the processing of the CMC expression data, we removed samples with RIN < 5.5. All samples in the dataset had read numbers exceeding 70 million. Reads had been normalized and transferred to log2-Counts per million (CPM) mapped reads using the voom function in limma (https://www.rdocumentation.org/packages/limma/versions/3.28.14). Genes with less than 1 CPM for more than half of the samples were considered not expressed and removed as in the CMC. To identify outliers, we converted raw reads to Fragments Per Kilobase of transcript per Million mapped reads (FPKM) and used hierarchical clustering to identify any sample(s) that clustered separately from the rest. We identified and removed 33 outlier samples. The R package Supervised Normalization of Microarrays (SNM) [Mecham and others 2010] was used to adjust by known (ancestry, sex, age at death, PMI, RIN & RIN squared) and hidden covariates, generated by surrogate variable analysis (as implemented in http://bioconductor.org/packages/release/bioc/html/sva.html). To be consistent with the CMC dataset, we excluded samples with age <17 yr. We also excluded SNPs absent or reported to have different alleles than listed on the PGC file. Ambiguous SNPs were also removed. Finally, 9386 SNPs, 9401 genes, 151 cases and 194 controls were included in the eQTL analysis. The cases included 83 Caucasian and 68 African American, and the controls 86 Caucasians, 108 African Americans.

#### eQTL analysis

We defined as cis-eQTL analysis the analysis of SNP - gene pairs closer than 500 KB and as trans- that of pairs at greater distance. In the trans-eQTL analysis we included all expressed genes but only SNPs in the schizophrenia loci. We performed all eQTL analyses with the matrixEQTL package using linear models [Shabalin 2012].

We performed eQTL analysis to select SNP-gene pairs for mediation analysis. Because we recognize that the top SZ associated SNP is not necessarily the one best capturing the effect on expression which may involve more than one variant, relatively loose thresholds were used to select SNP-gene pairs for mediation tests. For cis- and trans-eQTL, p < 0.01 and p < 10^-7 were required for proceeding to the next steps respectively. Our analysis showed that with these P value thresholds, we achieved FDR near 5% in both cases (CMC: cis 6.0%, trans 3.7%; BSC: cis 8.7%, trans 6.0%). Both cis- and trans- SNP-gene pairs that passed the thresholds were included in the mediation analysis.

#### Mediation analysis

Only significant SNP-gene pairs in the eQTL analysis as defined above were included in the mediation analysis. The Python package PyProcessMacro a public version of a widely used commercial mediation analysis tool PROCESS macro [Hayes 2018] (https://github.com/QuentinAndre/pyprocessmacro) performing a two-step linear regression model was applied considering one mediator at a time. The 1^st^ regression step was: M = ß_1_X + b + e; the 2^nd^ regression was: Y = ß_2_X + ß_3_M + b + e (were X: genotype, M: Mediator Gene expression, Y: Phenotype, ß: corresponding Coefficients, b: Intercept, e: Error). ß_2_ is the direct effect of the SNP on the phenotype. The indirect effect of a SNP on the phenotype through the mediator is the product of ß_1_ and ß_3_. The total effect is the sum of the indirect and the direct effects. PyProcessMacro calculates and provides 95% and 99% CI for the indirect (mediated) effect by 5000 bootstraps to test its statistical significance.

#### Permutations

To test the mediation significance accounting for our multiple testing we permuted the case/control assignment and repeated the same mediation analysis thus preserving the link between genotype and expression as well as the LD structure. We performed 100 permutations. For each permutation, we calculated the number of genes significant for mediation at the 95% and 99% CI. We then used these 100 values to calculate the mean number of genes positive for mediation under the null hypothesis and its standard deviation, from which we calculated the false discovery rate for the 95% and 99% CI based on the observed number of significant results.

## RESULTS

To identify our target pairs of gene-SNP group (set of SNPs in LD, see methods) for mediation analysis we first performed an eQTL analysis in the CMC and BSC datasets. The complete results are <u>reported in detail in Supplementary Table 2</u>. Below we provide a few metrics and highlight some of these results

### CMC dataset

In eQTL analysis in the CMC dataset we found 14,258 cis and 411 trans significant SNP-gene pairs (cis-: P < 0.01, trans-: P < 10^−7^, FDR ~5% for both). Of the 219 independent SNP groups (groups of SNPs in LD – see methods) 106 were significantly correlated with the expression of 311 genes in cis and 12 with 18 genes in trans. Of the 106 cis-eQTLs groups, 55 (51.9%) were correlated with the expression of more than one gene with a maximum of 22 for SNP group 80. Of the 311 genes with cis-eQTLs, 13 (4.2%) were correlated with 2 SNP groups at the same locus (<500 Kb). For trans-eQTLs, 3 out of 12 groups and their target genes were located on different chromosomes. These were group 19 on Chr1, which was an eQTL for *HECW1* (ENSG00000002746) on Chr7; Group 24 on Chr1 was an eQTL for *CDC27* (ENSG00000004897) on Chr17. Group 166 on Chr14, was an eQTL for genes *EPHA10* (ENSG00000183317) on Chr1 and *SLC35B1* (ENSG00000121073) on Chr17 Group166 was also a cis-eQTL for genes AC005477.1 and *RGS6*; genes *EPHA10* and *SLC35B1* had no nearby SZ-associated SNPs so it was not tested for cis-eQTL. Group 24 was a cis-eQTL for *FANCL*. *HECW1* was also not tested for cis-eQTL. The remaining 9 trans-eQTL groups were <6Mb away from the correlated genes and 7 of them were <1Mb. Four out of the 12 trans-eQTL SNP groups (33.3%) were correlated with more than one gene. Only 1 of these 4 groups (Group166) was located on a different chromosome from the correlated genes.

### BSC dataset

In eQTL analysis in the BSC dataset we found 8,958 cis and 220 trans significant SNP-gene pairs (cis-: P < 0.01, trans-: P < 10^−7^, FDR ~5%). Of the 219 independent SNP groups 92 were significantly correlated with the expression of 272 genes in cis and 7 with 7 genes in trans. Of the 92 cis-eQTLs groups, 52 (56.5%) were correlated with the expression of more than one gene with a maximum of 14 for SNP group 52. Of the 272 genes with cis-eQTL, 4 (1.5%) were correlated with 2 SNP groups at the same locus (<500 Kb). For trans-eQTL, 4 out of 7 groups and their target genes were located on different chromosomes. These were Group51 on Chr3 with gene *PSENEN* (ENSG00000205155) on Chr19 (Group51 was also cis-eQTL for gene *LRRFIP2*, *DCLK3, TRANK1*, while *PSENEN* did not have nearby SNP groups); Group11 on Chr1 with *OSBP* (ENSG00000110048) on Chr11 (Group11 was also cis-eQTL for *BRINP2*, while *OSBP* did not have nearby SNP groups); Group174 on Chr15 with *GMPS* (ENSG00000163655) on Chr3, (Group 174 was also cis-eQTL for PSMA4, *CRABP1, CTSH* while *GMPS* did not have nearby SNP groups); Group129 on Chr10, with *PPAPDC1B* (ENSG00000147535) on Chr8 (Group 129 was also cis-eQTL for genes *CNNM2*, C10orf32, *NT5C2*, RP11-18I14.10, *CALHM2*, *PSD*, *CUEDC2*, *PCGF6*, *FBXL15*, *TRIM8*, *TMEM180*, *INA*, while *PPAPDC1B* although it was within SNP group 111 did not have cis-eQTLs). All of the other groups were located in a <3 Mb region from the genes and 1 of them <1Mb.

### eQTL overlaps between CMC and BSC

Of the identified cis- and trans-eQTLs, 70 SNP groups that were eQTLs for 149 genes in cis, and 2 SNP groups that were eQTLs for 2 genes in trans respectively overlapped between the CMC and BSC datasets. Most of the cis-overlaps (68 of 70 SNP groups that were eQTLs for 137 genes) and all of the trans-overlaps showed consistent direction between the two datasets (consistent-only trans eQTL overlap - observed vs. expected Chi test = 1.2 x 10^−10^, data in Supplementary Table 2). Supplementary Figure 1 shows a scatterplot of the eQTL effect sizes observed in the two datasets at the individual SNP group and at the SNP level.

### Permutations to calculate false discovery (FDR)

As we discuss in the methods, the analysis platform we use to test mediation reports bootstrap-based confidence intervals (CI). Testing all correlated SNPs in each LD group complicates assessing the true significance of positive results. To correctly assess mediation significance and calculate reliable FDR we permuted the link between the genotypes and the phenotype (preserving only genotype-expression correlations), and repeated the mediation analysis, counting the number of positive results and comparing with the observed results.

For the CMC dataset, these permutations showed an average of 37.9 significant mediating effects with SD=7.4 for the 95% CI. Considering 113 observed mediating effects significant at 95% CI (see below), the FDR was 34%. Under the 99% CI, the mean number of genes with significant mediating effects was 9.54 (SD = 3.89). Considering 58 observed mediating effects significant at 99% CI, the FDR was 16.4%.

For the BSC dataset the mean of the number of genes with significant mediating effects in permutations was 32.16 (SD= 8.50); considering 178 observed positives (see below) the FDR was around 18.1%. Under the 99% CI, the mean of number of genes with significant mediating effects was 7.12 (SD= 3.31); considering 142 observed mediating effects significant at 99% CI, the FDR is 5.01%. More specific details are given on Table 1

**Table 1:** Results based on our data and on permutations for calculating FDR

|  | CMC | BSC |
| --- | --- | --- |
| Cases/Controls | 258/279 | 151/194 |
| SNPs | 9536 | 9386 |
| Genes | 16,311 | 9401 |
| Cis/Trans eQTL, 5% FDR | 14,258/411 | 8958/220 |
| 95% CI, observed | 113 | 178 |
| 95% CI, permutations | 37.9 ± 7.4 | 32.2 ± 8.5 |
| 95% CI, FDR | 34% | 18% |
| 99% CI, observed | 58 | 142 |
| 99% CI, permutations | 9.5 ± 3.9 | 7.1 ± 3.3 |
| 99% CI, FDR | 16.4% | 5.0% |

### CMC Mediation results

In the CMC dataset, 4,156 of the 14,669 SNP-gene pairs were significant for mediation (the 95% CI calculated by bootstrapping did not include 0, see methods). This corresponds to 68 SNP groups (of the 106 tested - 64%) significantly mediated by 113 genes under the 95% CI model (Supplementary Table 3). About half (56) of the 113 genes showed a negative mediating effect, meaning that the direction in which the risk allele changed expression was protecting against the disease. Under the 99% CI model, the number of significant SNP-gene pairs was 1798 which consisted of 40 SNP groups that were eQTLs for 58 genes. 30 of these genes showed a negative mediating effect.

### BSC Mediation results

In the BSC dataset, 5,575 out of 9,178 pairs were significant for the mediation, corresponding to 78 of the 92 tested SNP groups (85%) significantly mediated by 178 genes under 95% CI model (Supplementary Table 3). Similar to CMC, about half (93) of the 178 genes (52.9%) showed a negative mediating effect. Under the 99% CI model, the number of significant SNP-gene pairs was 4340 consisting of 67 SNP groups and 142 genes. 77 of these 142 genes showed a negative mediating effect.

### Mediation overlaps between CMC and BSC

Supplementary Table 3 shows the mediation results and for each mediation that was significant at least in one dataset it also shows the corresponding effects of gene expression on disease. The vast majority of mediation positives at the 95% confidence level also showed significant effects of gene expression on disease at p<0.05 (BSC: 161/180, 89.4%; CMC: 77/114, 67.5%). That was at or near 100% for mediation positives at the 99% confidence level (BSC: 143/143, 100%; CMC: 57/58, 98%). Supplementary Table 3 also shows SNPs within each group that were present in both datasets along with their mediation effect when it is significant. When were more than one such SNPs overlap the one with lowest SZ GWAS p-value is shown.

To validate the direction of mediation across datasets we examined the overlapping SNP group – gene pairs between datasets for consistency. Of the 153 genes included in both datasets mediation analysis 38 were significant in both, under the 95% CI model. Of those 38, 33 (87%) showed consistent direction in the two studies (Chi^2^ test P=6*10^−6^). Of these 33, 18 had positive and 15 had negative mediating effect. Under the 99% CI model, 20 genes showed significant mediating effect in both datasets 18 of them (90%, Chi2 test P=3.47*10^−4^)) with consistent directions. Eleven of them (*<u>SOWAHC, AKAP6, GOLGA2P7, GPN1, FTSJ2, NDUFA2, THOC7, NAGA, CSPG4P12, DDHD2, RERE</u>*) had positive and seven (*<u>MAU2, TYW5, TDRD9, ZMAT2, CNTN4, SF3B1, NISCH</u>*) had negative effects as defined above. Supplementary Figure 2 shows a scatterplot of the mediation effect sizes observed in the two datasets at the SNP group and at the individual SNP level.

### Further validations

To ensure our results are not driven by SNPs that are not truly associated with SZ we removed all SNPs with GWAS p>10^−8^ and reassessed the results. We note that this is likely to also remove many true positives, not because many of these SNPs that are also eQTL might not represent the strongest signal but might contribute to it along with others at the same locus, the lead SNP being only the proxy of multiple functional variants. In the CMC 49 genes remained significant at 95% CI and 25 under the 99% CI. In the BSC these numbers were 72 and 62 respectively. All FDRs were similar. The overlaps were 18 and 8 respectively and their 94.4% and 100% concordant. Among these concordant mediation results 47.1% and 37.5% were negative, still supporting that the direction in which expression affects risk is often different from the direction in which the risk allele changes expression.

To assess whether stricter thresholds for eQTL would increase the fraction of consistent and exclude false positive eQTLs as a confounder, in addition to cis thresholds of 10^−2^, we also performed mediation analysis using cis eQTL threshold 10^−4^. The overlaps were 16 and 12 for the 95 and 99% CI respectively and the directions of all but one was concordant in both cases. Among these concordant mediation results 7/15 and 4/11 were negative.

Further, to ensure out results are not due to confounding due to stratification despite our correction for ethnicity, we repeated the analysis of previously significant mediation results using only the larger Caucasian subset of the BSC. In this subset 91 genes remained significant for mediation, 41 positive and 51 negative, excluding race as a possible confounder of the directional effect.

## DISCUSSION

In view of the fact that SNPs are often eQTLs for multiple genes, many of these genes are likely unknown, our goal was to test whether genetic associations between common variants and disease are mediated by gene regulation and to determine whether the direction of this mediation is that expected from the direction in which the risk allele modifies the expression. To achieve this goal we performed a mediation analysis on the two largest independent gene expression datasets from the dorsolateral prefrontal cortex of SZ cases and controls - the CMC and the BSC. We tested variants associated with schizophrenia that are also eQTL in these datasets. Due to the complexities of combining GWAS data and expression data from different datasets in the presence of LD, where different SNPs can represent the same association/eQTL signal, we report on SNP groups, defined as SNPs connected by LD. To avoid false positives due to the multiple testing within and between these groups, we defined nominal significance thresholds based on confidence intervals and then calculated FDR based on permutations.

Transcriptome-wide association studies (TWAS) test a hypothesis similar to mediation and may be subject similar challenges in interpretation as well. TWAS use expression and SNP data to estimate the genetic component of gene expression, which is then applied to GWAS data to test for a genetical gene expression difference between cases and controls. While an observed effect is typically attributed to the expression level of the gene in question, the same SNPs may influence the expression levels of other genes. The genetic effect is then the sum over all the genes influenced, rather than just the gene nominally considered by the TWAS.

Our full list of eQTL is provided on Supplementary Table 2 and as expected, is similar to previous eQTL reported for these same datasets [Fromer and others 2016; Jaffe and others 2018]. By reducing the search space to only SZ-associated SNPs, we also identified trans-eQTL described above. Many of the trans-eQTL were also cis-eQTL affecting multiple genes, and many of the genes affected in trans were at locations that showed no genetic associations with SZ. It is possible that these are true risk genes but they are not implicated by genetics either because of an absence of local regulatory variation or because local regulatory variation influences other genes in discordant directions and the sum of effects on risk is minimal.

We find that a large number of GWAS signals show evidence of mediation by at least one of the genes for which they that are eQTL, as is generally accepted by the literature. Mediation evidence is found for 64% of SNP groups in the CMC dataset and 85% in the BSC dataset at the relaxed criteria FDR of 33.5% and 18.1% respectively. This suggests that more than half of the signals are mediated by gene expression. Given power limitations, this is likely a low estimate.

Twenty genes showed significant mediating effect in both datasets under our 99% CI model with FDR 16% and and 5% in the CMC BSC respectively, 18 of them with conserved directions: *SOWAHC, AKAP6, GOLGA2P7, GPN1, FTSJ2, NDUFA2, THOC7, NAGA, CSPG4P12, DDHD2, RERE* with same-direction genetic and gene expression directions, and *MAU2, TYW5, TDRD9, ZMAT2, CNTN4, SF3B1, NISCH* with opposite-direction genetic and gene expression directions. We consider these high confidence schizophrenia genes.

SNP group 52 is an interesting example. This group on chromosome 3 52-52 Mb regulates 20 genes across the two DLPFC datasets, 11 in both, of which 3 are significant for mediation (*GNL3, PPM1M*, and *NISCH*) in both datasets. The GTEx database shows the lead SNP of this group to be eQTL for 23 genes across tissues, mostly overlapping with these 20 genes, of which two are in the cerebral hemispheres (*GNL3, PPM1M*). For *PPM1M* the mediation is in the direction suggested by the risk allele effect, but for *GNL3* and *NISCH* it is negative. None of the 3 genes reaches significance in a recent TWAS [Hall and others 2020], where a different 4 genes do so in the region *NEK4, TMEM110, GLT8D1, ITIH4*. The first three are also eQTL in our results but none shows mediation.

We identify multiple instances where the mediation is not in the direction suggested by the effect of the risk allele on expression. This means that although the disease risk allele correlates with decreased expression of a gene, the decrease in expression is associated with lower risk, or alternatively that the risk allele increases expression but increased expression associates with lower risk. The validity of this result is supported not only by its presence in both datasets, but also by the high consistency of the overlapping signals. Further, this result holds when we apply more stringent thresholds for GWAS significance and eQTL significance and when we restrict the analysis to Caucasians. This observation is of great importance for the design of studies of the biological link between genetic variation and disease. For example, examining the consequences of a gene knock down under the wrong assumption that lower expression mediated higher risk could lead to incorrect or misleading conclusions. We hypothesize that this commonly observed apparent discordance in direction is because it is common for variants to regulate multiple genes. This one-to-many relationship in our results could be a general phenomenon if one accounts for statistical power and the study of mixed cell populations at single time points. Given the strong selective disadvantage of schizophrenia [Power and others 2013], variants whose effects counteract each other may be more likely to persist in a population. In this case, the small effect sizes of GWAS variants on disease risk may reflect their net effect on multiple genes, with the individual effects possibly larger. Another interesting way forward for future analyses would be to model the aggregate effect of many SNPs on a gene’s expression and to examine mediation under this model as this could further elucidate the nature of the SNP → gene → SZ relationship.

Our results highlight the complexity of the interplay between population genetics and regulatory variation, which creates unpredictable relationships between the effects of variants of gene expression and that of gene expression on the risk. As it becomes increasingly common to manipulate the genome in targeted ways in order to understand the biology basis of disease risk represented by GWAS variants [Das and others 2020], understanding this interplay is increasingly important. At the same time however, this opens the possibility that small observed effects of variants on risk might underestimate the effects of individual gene expression levels on the risk, which in turn could open new possibilities for therapeutic interventions.

## DATA ACCESS

All data used in this work was already publicly available. No new data was generated.

## ACKNOWLEDGEMENTS

This work was supported by NIH grant R01MH113215. The corresponding author is also supported by P50MH094268, R01MH106522 and RF1MH122936.

## DISCLOSURE DECLARATIONS

The authors have no conflicts of interest or other disclosures

## Notes

### Competing Interest Statement

The authors have declared no competing interest.

### Summary of Updates

Revised manuscript, supplementary tables and new supplementary figure

## BIBLIOGRAPHY

Akerborg O, Spalinskas R, Pradhananga S, Anil A, Hojer P, Poujade FA, Folkersen L, Eriksson PP, Sahlen P. 2019. High-Resolution Regulatory Maps Connect Vascular Risk Variants to Disease-Related Pathways. Circ Genom Precis Med 12(3):e002353.

Avramopoulos D. 2018. Recent Advances in the Genetics of Schizophrenia. Mol Neuropsychiatry 4(1):35–51.

Cardno AG, Marshall EJ, Coid B, Macdonald AM, Ribchester TR, Davies NJ, Venturi P, Jones LA, Lewis SW, Sham PC, Gottesman, II, Farmer AE, McGuffin P, Reveley AM, Murray RM. 1999. Heritability estimates for psychotic disorders: the Maudsley twin psychosis series. Arch Gen Psychiatry 56(2):162–168.

Consortium SPG-WASG. 2011. Genome-wide association study identifies five new schizophrenia loci. Nat Genet 43(10):969–976.

Consortium SWGotPG. 2014. Biological insights from 108 schizophrenia-associated genetic loci. Nature 511(7510):421–427.

Cookson W, Liang L, Abecasis G, Moffatt M, Lathrop M. 2009. Mapping complex disease traits with global gene expression. Nat Rev Genet 10(3):184–194.

Das D, Feuer K, Wahbeh M, Avramopoulos D. 2020. Modeling Psychiatric Disorder Biology with Stem Cells. Curr Psychiatry Rep 22(5):24.

Fromer M, Roussos P, Sieberts SK, Johnson JS, Kavanagh DH, Perumal TM, Ruderfer DM, Oh EC, Topol A, Shah HR, Klei LL, Kramer R, Pinto D, Gumus ZH, Cicek AE, Dang KK, Browne A, Lu C, Xie L, Readhead B, Stahl EA, Xiao J, Parvizi M, Hamamsy T, Fullard JF, Wang YC, Mahajan MC, Derry JM, Dudley JT, Hemby SE, Logsdon BA, Talbot K, Raj T, Bennett DA, De Jager PL, Zhu J, Zhang B, Sullivan PF, Chess A, Purcell SM, Shinobu LA, Mangravite LM, Toyoshiba H, Gur RE, Hahn CG, Lewis DA, Haroutunian V, Peters MA, Lipska BK, Buxbaum JD, Schadt EE, Hirai K, Roeder K, Brennand KJ, Katsanis N, Domenici E, Devlin B, Sklar P. 2016. Gene expression elucidates functional impact of polygenic risk for schizophrenia. Nat Neurosci 19(11):1442–1453.

Gilad Y, Rifkin SA, Pritchard JK. 2008. Revealing the architecture of gene regulation: the promise of eQTL studies. Trends Genet 24(8):408–415.

Hall LS, Medway CW, Pain O, Pardinas AF, Rees EG, Escott-Price V, Pocklington A, Bray NJ, Holmans PA, Walters JTR, Owen MJ, O’Donovan MC. 2020. A transcriptome-wide association study implicates specific pre- and post-synaptic abnormalities in schizophrenia. Hum Mol Genet 29(1):159–167.

Hayes AF. 2018. Introduction to mediation, moderation, and conditional process analysis : a regression-based approach. New York: Guilford Press. xx, 692 pages p.

Hill MJ, Jeffries AR, Dobson RJ, Price J, Bray NJ. 2012. Knockdown of the psychosis susceptibility gene ZNF804A alters expression of genes involved in cell adhesion. Hum Mol Genet 21(5):1018–1024.

Hindorff LA, Sethupathy P, Junkins HA, Ramos EM, Mehta JP, Collins FS, Manolio TA. 2009. Potential etiologic and functional implications of genome-wide association loci for human diseases and traits. Proc Natl Acad Sci U S A 106(23):9362–9367.

Jaffe AE, Straub RE, Shin JH, Tao R, Gao Y, Collado-Torres L, Kam-Thong T, Xi HS, Quan J, Chen Q, Colantuoni C, Ulrich WS, Maher BJ, Deep-Soboslay A, Cross AJ, Brandon NJ, Leek JT, Hyde TM, Kleinman JE, Weinberger DR. 2018. Developmental and genetic regulation of the human cortex transcriptome illuminate schizophrenia pathogenesis. Nat Neurosci 21(8):1117–1125.

Kety SS. 1987. The significance of genetic factors in the etiology of schizophrenia: results from the national study of adoptees in Denmark. J Psychiatr Res 21(4):423–429.

Mackinnon DP, Fairchild AJ. 2009. Current Directions in Mediation Analysis. Curr Dir Psychol Sci 18(1):16.

MacKinnon DP, Fairchild AJ, Fritz MS. 2007. Mediation analysis. Annu Rev Psychol 58:593–614.

Majewski J, Pastinen T. 2011. The study of eQTL variations by RNA-seq: from SNPs to phenotypes. Trends Genet 27(2):72–79.

Manolio TA, Collins FS, Cox NJ, Goldstein DB, Hindorff LA, Hunter DJ, McCarthy MI, Ramos EM, Cardon LR, Chakravarti A, Cho JH, Guttmacher AE, Kong A, Kruglyak L, Mardis E, Rotimi CN, Slatkin M, Valle D, Whittemore AS, Boehnke M, Clark AG, Eichler EE, Gibson G, Haines JL, Mackay TF, McCarroll SA, Visscher PM. 2009. Finding the missing heritability of complex diseases. Nature 461(7265):747–753.

Maurano MT, Humbert R, Rynes E, Thurman RE, Haugen E, Wang H, Reynolds AP, Sandstrom R, Qu H, Brody J, Shafer A, Neri F, Lee K, Kutyavin T, Stehling-Sun S, Johnson AK, Canfield TK, Giste E, Diegel M, Bates D, Hansen RS, Neph S, Sabo PJ, Heimfeld S, Raubitschek A, Ziegler S, Cotsapas C, Sotoodehnia N, Glass I, Sunyaev SR, Kaul R, Stamatoyannopoulos JA. 2012. Systematic localization of common disease-associated variation in regulatory DNA. Science 337(6099):1190–1195.

Mecham BH, Nelson PS, Storey JD. 2010. Supervised normalization of microarrays. Bioinformatics 26(10):1308–1315.

Messias EL, Chen CY, Eaton WW. 2007. Epidemiology of schizophrenia: review of findings and myths. Psychiatr Clin North Am 30(3):323–338.

O’Brien HE, Hannon E, Hill MJ, Toste CC, Robertson MJ, Morgan JE, McLaughlin G, Lewis CM, Schalkwyk LC, Hall LS, Pardinas AF, Owen MJ, O’Donovan MC, Mill J, Bray NJ. 2018. Expression quantitative trait loci in the developing human brain and their enrichment in neuropsychiatric disorders. Genome Biol 19(1):194.

Pardinas AF, Holmans P, Pocklington AJ, Escott-Price V, Ripke S, Carrera N, Legge SE, Bishop S, Cameron D, Hamshere ML, Han J, Hubbard L, Lynham A, Mantripragada K, Rees E, MacCabe JH, McCarroll SA, Baune BT, Breen G, Byrne EM, Dannlowski U, Eley TC, Hayward C, Martin NG, McIntosh AM, Plomin R, Porteous DJ, Wray NR, Caballero A, Geschwind DH, Huckins LM, Ruderfer DM, Santiago E, Sklar P, Stahl EA, Won H, Agerbo E, Als TD, Andreassen OA, Baekvad-Hansen M, Mortensen PB, Pedersen CB, Borglum AD, Bybjerg-Grauholm J, Djurovic S, Durmishi N, Pedersen MG, Golimbet V, Grove J, Hougaard DM, Mattheisen M, Molden E, Mors O, Nordentoft M, Pejovic-Milovancevic M, Sigurdsson E, Silagadze T, Hansen CS, Stefansson K, Stefansson H, Steinberg S, Tosato S, Werge T, Collier DA, Rujescu D, Kirov G, Owen MJ, O’Donovan MC, Walters JTR. 2018. Common schizophrenia alleles are enriched in mutation-intolerant genes and in regions under strong background selection. Nat Genet 50(3):381–389.

Power RA, Kyaga S, Uher R, MacCabe JH, Langstrom N, Landen M, McGuffin P, Lewis CM, Lichtenstein P, Svensson AC. 2013. Fecundity of patients with schizophrenia, autism, bipolar disorder, depression, anorexia nervosa, or substance abuse vs their unaffected siblings. JAMA Psychiatry 70(1):22–30.

Ripke S, O’Dushlaine C, Chambert K, Moran JL, Kahler AK, Akterin S, Bergen SE, Collins AL, Crowley JJ, Fromer M, Kim Y, Lee SH, Magnusson PK, Sanchez N, Stahl EA, Williams S, Wray NR, Xia K, Bettella F, Borglum AD, Bulik-Sullivan BK, Cormican P, Craddock N, de Leeuw C, Durmishi N, Gill M, Golimbet V, Hamshere ML, Holmans P, Hougaard DM, Kendler KS, Lin K, Morris DW, Mors O, Mortensen PB, Neale BM, O’Neill FA, Owen MJ, Milovancevic MP, Posthuma D, Powell J, Richards AL, Riley BP, Ruderfer D, Rujescu D, Sigurdsson E, Silagadze T, Smit AB, Stefansson H, Steinberg S, Suvisaari J, Tosato S, Verhage M, Walters JT, Levinson DF, Gejman PV, Kendler KS, Laurent C, Mowry BJ, O’Donovan MC, Owen MJ, Pulver AE, Riley BP, Schwab SG, Wildenauer DB, Dudbridge F, Holmans P, Shi J, Albus M, Alexander M, Campion D, Cohen D, Dikeos D, Duan J, Eichhammer P, Godard S, Hansen M, Lerer FB, Liang KY, Maier W, Mallet J, Nertney DA, Nestadt G, Norton N, O’Neill FA, Papadimitriou GN, Ribble R, Sanders AR, Silverman JM, Walsh D, Williams NM, Wormley B, Arranz MJ, Bakker S, Bender S, Bramon E, Collier D, Crespo-Facorro B, Hall J, Iyegbe C, Jablensky A, Kahn RS, Kalaydjieva L, Lawrie S, Lewis CM, Lin K, Linszen DH, Mata I, McIntosh A, Murray RM, Ophoff RA, Powell J, Rujescu D, Van Os J, Walshe M, Weisbrod M, Wiersma D, Donnelly P, Barroso I, Blackwell JM, Bramon E, Brown MA, Casas JP, Corvin AP, Deloukas P, Duncanson A, Jankowski J, Markus HS, Mathew CG, Palmer CN, Plomin R, Rautanen A, Sawcer SJ, Trembath RC, Viswanathan AC, Wood NW, Spencer CC, Band G, Bellenguez C, Freeman C, Hellenthal G, Giannoulatou E, Pirinen M, Pearson RD, Strange A, Su Z, Vukcevic D, Donnelly P, Langford C, Hunt SE, Edkins S, Gwilliam R, Blackburn H, Bumpstead SJ, Dronov S, Gillman M, Gray E, Hammond N, Jayakumar A, McCann OT, Liddle J, Potter SC, Ravindrarajah R, Ricketts M, Tashakkori-Ghanbaria A, Waller MJ, Weston P, Widaa S, Whittaker P, Barroso I, Deloukas P, Mathew CG, Blackwell JM, Brown MA, Corvin AP, McCarthy MI, Spencer CC, Bramon E, Corvin AP, O’Donovan MC, Stefansson K, Scolnick E, Purcell S, McCarroll SA, Sklar P, Hultman CM, Sullivan PF. 2013. Genome-wide association analysis identifies 13 new risk loci for schizophrenia. Nat Genet 45(10):1150–1159.

Schaub MA, Boyle AP, Kundaje A, Batzoglou S, Snyder M. 2012. Linking disease associations with regulatory information in the human genome. Genome Res 22(9):1748–1759.

Schrode N, Ho SM, Yamamuro K, Dobbyn A, Huckins L, Matos MR, Cheng E, Deans PJM, Flaherty E, Barretto N, Topol A, Alganem K, Abadali S, Gregory J, Hoelzli E, Phatnani H, Singh V, Girish D, Aronow B, McCullumsmith R, Hoffman GE, Stahl EA, Morishita H, Sklar P, Brennand KJ. 2019. Synergistic effects of common schizophrenia risk variants. Nat Genet 51(10):1475–1485.

Shabalin AA. 2012. Matrix eQTL: ultra fast eQTL analysis via large matrix operations. Bioinformatics 28(10):1353–1358.

Visel A, Rubin EM, Pennacchio LA. 2009. Genomic views of distant-acting enhancers. Nature 461(7261):199–205.

Yang CP, Li X, Wu Y, Shen Q, Zeng Y, Xiong Q, Wei M, Chen C, Liu J, Huo Y, Li K, Xue G, Yao YG, Zhang C, Li M, Chen Y, Luo XJ. 2018. Comprehensive integrative analyses identify GLT8D1 and CSNK2B as schizophrenia risk genes. Nat Commun 9(1):838.

Zhao X, Lynch J, xa, G, Chen Q, John Deighton served as e, Gavan Fitzsimons served as associate editor for this a. 2010. Reconsidering Baron and Kenny: Myths and Truths about Mediation Analysis. Journal of Consumer Research 37(2):197–206.

