## Supplementary figures for "Schizophrenia Risk Alleles Often Affect the Expression of Many Genes and Each Gene May Have a Different Effect on The Risk; A Mediation Analysis"

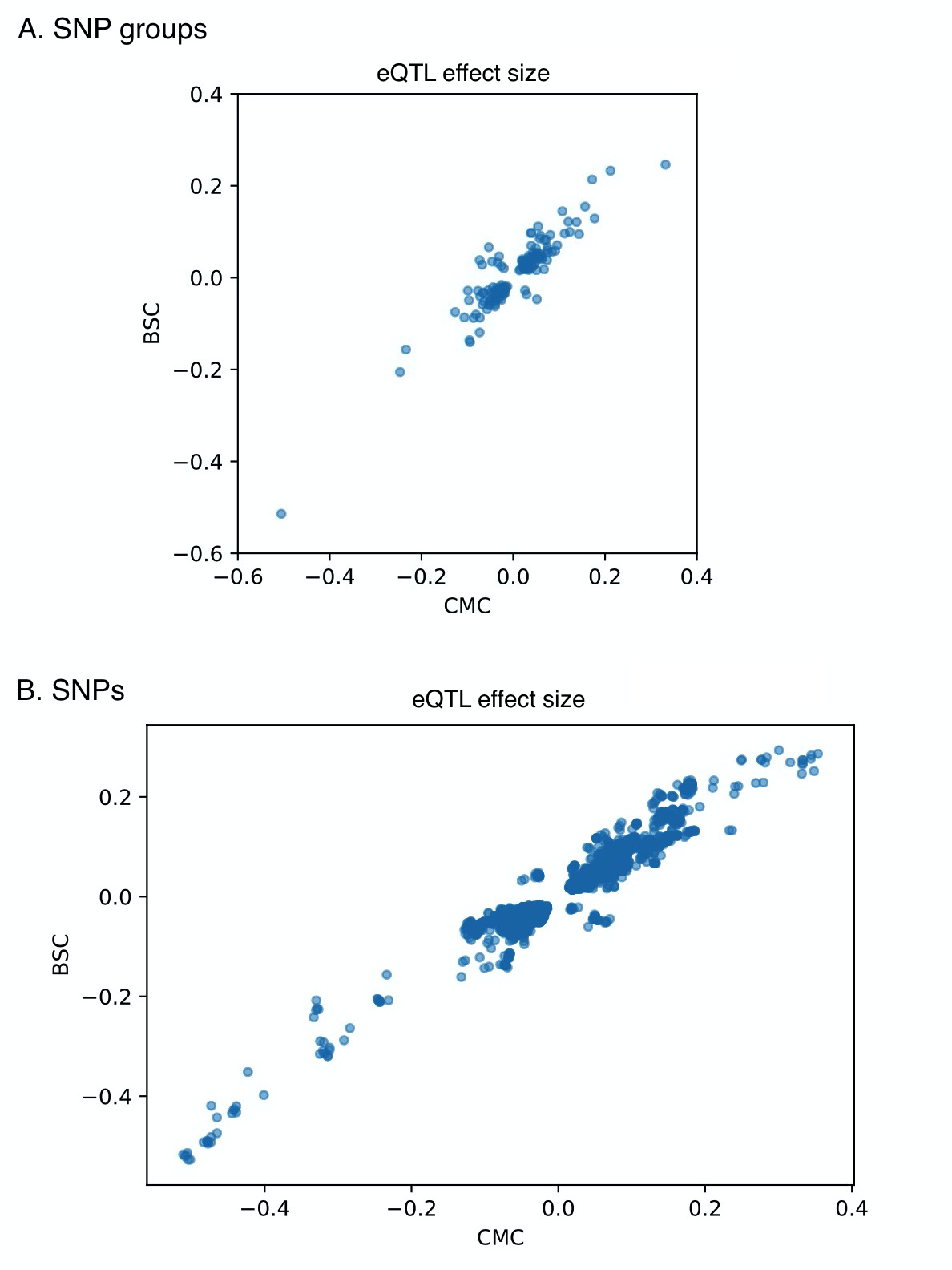

Supplementary Figure 1: Plot of the eQTL effect size in the CMC versus the BSC dataset at the SNP group level (A) and at the single SNP level (B)

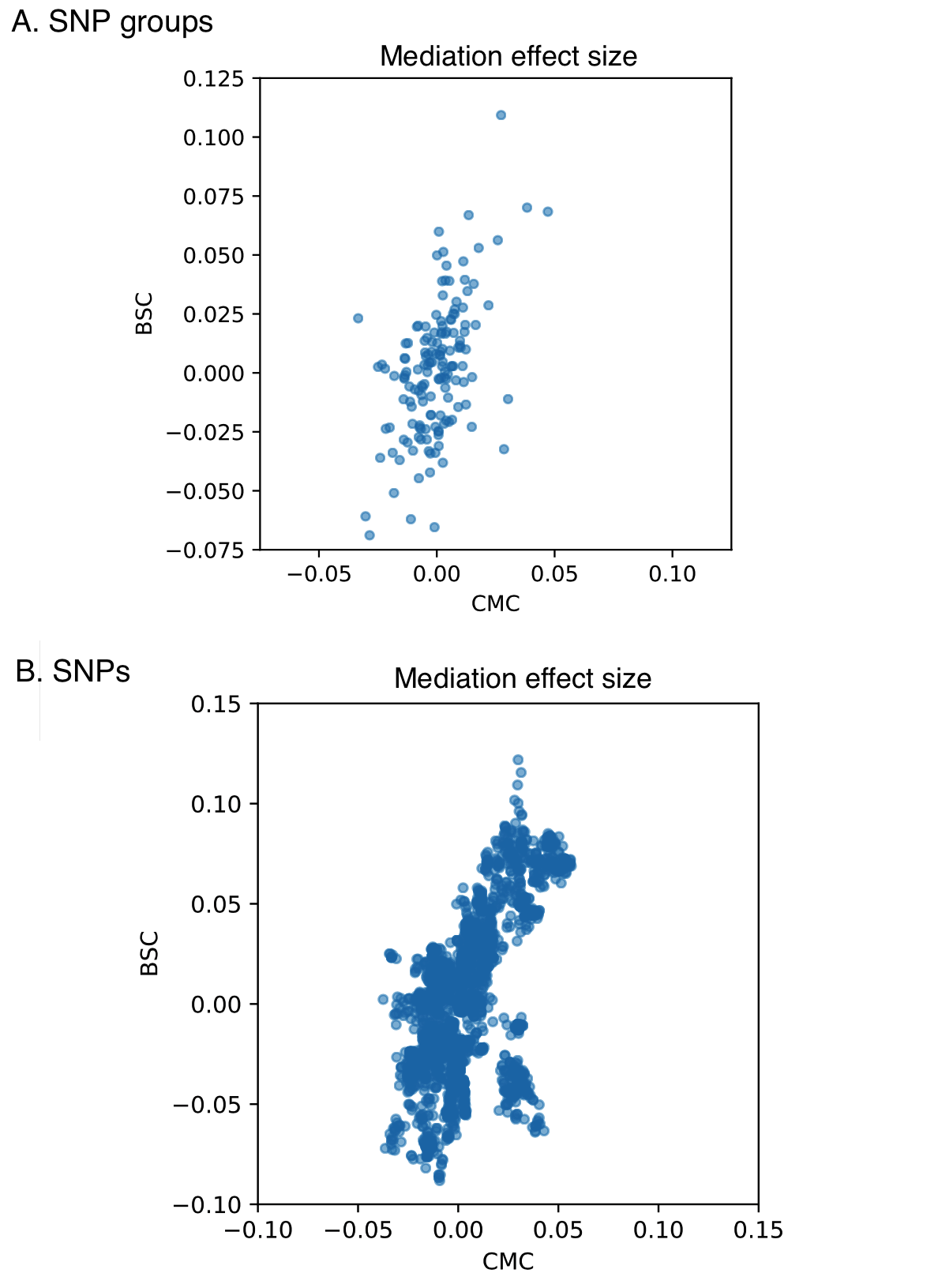

Supplementary Figure 2: Plot of the mediation effect size in the CMC versus the BSC dataset at the SNP group level (A) and at the single SNP level (B). The cluster on the lower right corresponds to two large SNP groups (129, 191) that include 280 SNPs and happen to be discordant. None of them is significant in both datasets.
